# Emerging patterns of plasmid-host coevolution that stabilize antibiotic resistance

**DOI:** 10.1101/146118

**Authors:** Thibault Stalder, Linda M. Rogers, Chris Renfrow, Hirokazu Yano, Zachary Smith, Eva M. Top

**Affiliations:** Department of Biological Sciences, University of Idaho, Moscow, Idaho, USA; Institute for Bioinformatics and Evolutionary Studies, University of Idaho, Moscow, Idaho, USA

**Keywords:** antibiotic resistance, plasmid, horizontal gene transfer, evolution, adaptation

## Abstract

Multidrug resistant bacterial pathogens have become a serious global human health threat, and conjugative plasmids are important drivers of the rapid spread of resistance to last-resort antibiotics. Whereas antibiotics have been shown to select for adaptation of resistance plasmids to their new bacterial hosts, or *vice versa*, a general evolutionary mechanism has not yet emerged. Here we conducted an experimental evolution study aimed at determining general patterns of plasmid-bacteria evolution. Specifically, we found that a large conjugative resistance plasmid follows the same evolutionary trajectories as its non-conjugative mini-replicon in the same and other species. Furthermore, within a single host–plasmid pair three distinct patterns of adaptive evolution led to increased plasmid persistence: i) mutations in the replication protein gene (*trfA1*); ii) the acquisition by the resistance plasmid of a transposon from a co-residing plasmid encoding a putative toxin-antitoxin system; iii) a mutation in the host’s global transcriptional regulator gene *fur*. Since each of these evolutionary solutions individually have been shown to increase plasmid persistence in other plasmid-host pairs, our work points towards common mechanisms of plasmid stabilization. These could become the targets of future alternative drug therapies to slow down the spread of antibiotic resistance.

## INTRODUCTION

The rapid evolution of multidrug resistant bacteria is a fundamental threat to human health that requires immediate action. ^1–3^. Acquisition of resistance to antibiotics by bacteria is largely caused by horizontal gene transfer, resulting in rapid adaptation of bacteria to almost all antibiotics ^4–6^. This DNA exchange is in part facilitated by conjugative plasmids ^7^, and can lead to antibiotic treatment failure ^8,9^. Equally disturbing is that plasmids are increasingly reported as vectors of spread of resistance to last line antibiotics such as colistin or carbapenems ^10,11^. Understanding the evolutionary trajectories that lead plasmids to spread and persist in bacterial communities is indispensable to developing efficient strategies to fight antibiotic resistance.

Although horizontally acquired antibiotic resistance genes and plasmids can confer a cost to the bacteria and are therefore expected to be lost again, antibiotic resistance is not always reversible; even after removal of the antibiotic selecting for resistance, antibiotic resistant bacteria can persist within bacterial populations ^12,13^. In the case of plasmids this phenomenon is often referred as the plasmid paradox: based on evolutionary principles, plasmids shouldn’t persist in bacterial populations, yet they do ^14–16^. On a more general level, plasmid-bacteria evolution can give us important clues about how symbiotic interactions evolve along a parasitism-mutualism continuum ^16,17^.

*In vitro* experimental evolution studies have become the strategy of choice to gain insight into how antibiotic therapy drives plasmid-host evolution and long-term persistence of antibiotic resistance plasmids. Such surveys have highlighted that compensatory mutations occurring during evolution can reduce the cost of carriage of the plasmid, change the horizontal transferability of the plasmid, or improve the vertical inheritance of the plasmid to daughter cells ^16,18–30^. More generally, these compensatory mutations result in increased persistence of the plasmids in bacterial populations or communities. Some of these studies showed that mutation(s) could occur either in the host chromosome only ^16,18,28,29^, on the plasmid only ^26,30^ or both ^19,22,23,29^. The most recent of these showed how specific mutations in some hosts could decrease the cost of carriage of the plasmid by restoring the general transcription pattern of the host, which was disturbed after acquisition of the plasmid ^16,31^. Other recent studies pointed out that plasmid evolution could lead to increased persistence of the plasmid by improving its vertical inheritance through acquisition of a transposon from a co-residing native plasmid containing a toxin/antitoxin system ^29^, or by removing an apparently costly plasmid region, encompassing transfer genes ^30^. All these surveys point to the diversity of evolutionary trajectories by which a plasmid-host adapt to each other, and how evolution shapes the trade-off between horizontal and vertical modes of transmission ^21^.

In one of our studies we showed that a mini-replicon called pMS0506 adapted to a novel host, *Shewanella oneidensis* MR-1 (here after designated as “MR-1”) within 1,000 generations of growth under selection for the plasmid ^26,27^. This non-conjugative plasmid was derived from the broad-host-range plasmid of the plasmid incompatibility group IncP-1β, pBP136 ^32^. The only evolutionary changes in the plasmid were mutations in the helicase-binding domain of the plasmid replication initiation protein TrfA1. These led to decreased interactions with host-encoded DnaB helicase, likely ameliorating the interference cost of the plasmid and thereby stabilizing the plasmid in the population ^26,33^.

Conjugation is an intrinsic part of the life cycle of conjugative plasmids and can affect how plasmids and novel bacterial hosts adapt to each other. Therefore, we set out to compare the evolutionary trajectories of the native conjugative plasmid with that of its previously studied non-conjugative mini-replicon. In this study we address the following three questions: Does experimental evolution of the native, conjugative plasmid pBP136 in the same host wherein its mini-replicon was evolved, also lead to improve plasmid persistence? If so, how do the evolutionary trajectories differ between the two plasmids? Do we begin to see common evolutionary solutions for the stabilization of incoming drug resistance plasmids? In combination with recently published findings on other plasmid-bacteria model systems our results point towards general mechanisms of stabilization of antibiotic resistance plasmids in various bacterial species.

## RESULTS

### Plasmid-host coevolution improves plasmid persistence

We first tested the hypothesis that the self-transmissible plasmid pBP136Km can become more persistent in strain MR-1 over time, just like its mini-replicon. Five replicate populations of MR-1 (pBP136Km) were evolved in serial batch culture under kanamycin selection, and after 1,000 generations three randomly chosen clones from each population were tested for plasmid persistence in the absence of antibiotics. While initially unstable in this host, pBP136Km showed significantly greater persistence in evolved clones from four of the five lineages, L1-L4. In each case, after a 6-day plasmid persistence assay the plasmid was still retained in at least 70% of the cells, whereas the ancestral plasmid was present in fewer than 2%. Detailed results are shown for clones from populations L1 and L4 (Fig. 1). The data confirm previous studies that persistence of conjugative plasmids can rapidly improve over evolutionary time.

**Figure 1.**
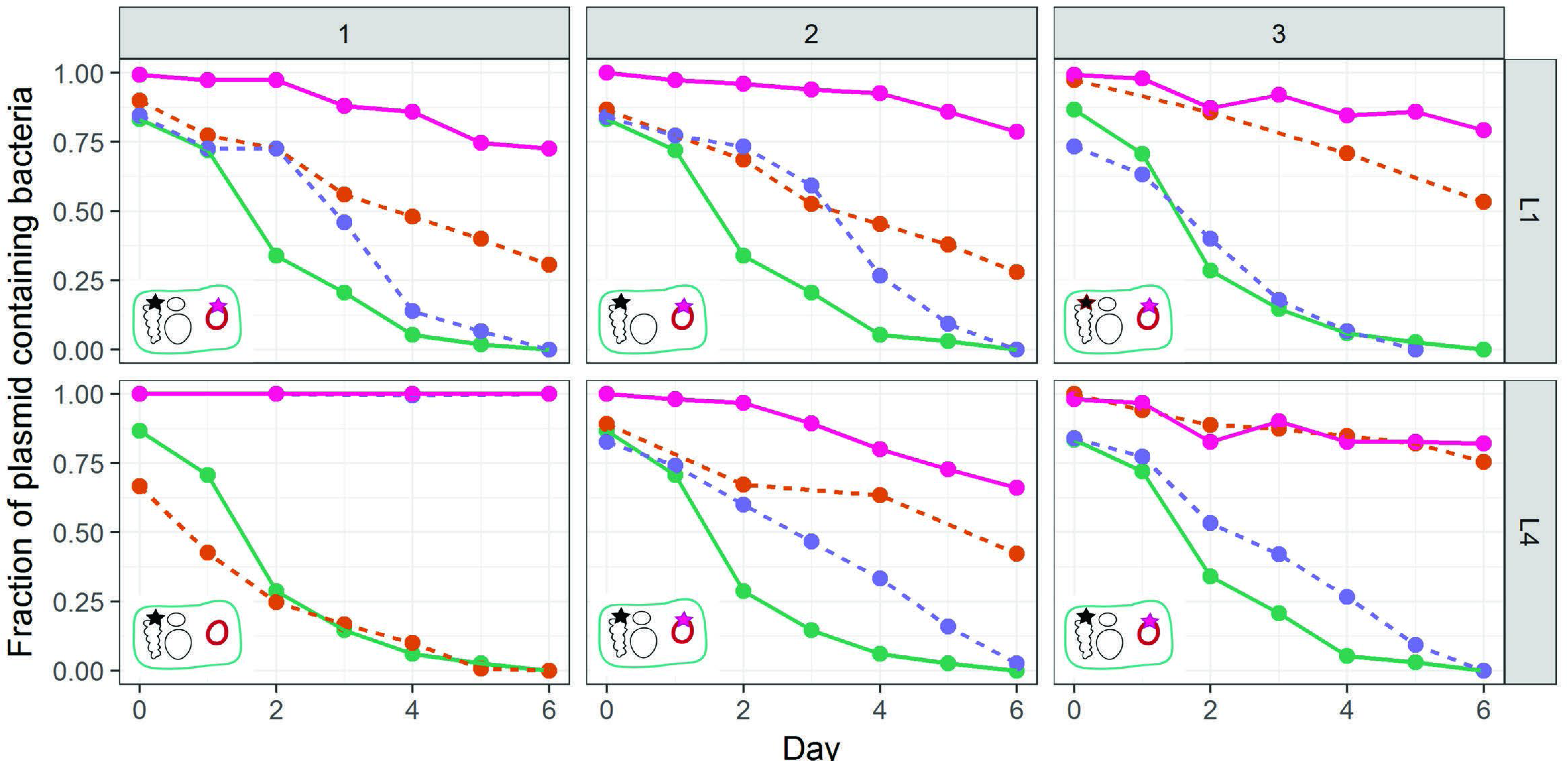
Persistence of the ancestral and evolved plasmids in the ancestral and co-evolved hosts from lineages L1 and L4 (clones L1-1, L1-2, L1-3 and L4-1, L4-2, L4-3, indicated as 1,2 and 3 above the panels). Solid lines: ancestral and evolved clones; dashed line: permuations of evolved/ancestral host/plasmids; 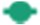: ancestral plasmid in ancestral host; 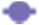 : ancestral plasmid in evolved host; 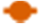: evolved plasmid in ancestral host, 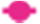 evolved plasmid in its co-evolved host. Each line is the mean of 3 individual replicates. For clarity error bars are not shown but the previously described plasmid population dynamics model ^29,34,35^ was used to determine if the persistence dynamics of two bacteria-plasmid pairs where similar or not (see Materials and Methods and SI). Inserts in the bottom left of each graph depict with a star the location of the mutation(s) (plasmid or chromosome) responsible for the increased plasmid persistence. Each replicon wihin the evolved cells are shown: the chromosome, the native plasmids (black circles) and the plasmid pBP136Km (pink circle). Note: plasmid-free segregants of evolved hosts were obtained after replicating the evolved plasmid-containing host onto non-selective media.

### Multiple evolutionary pathways can lead to improve plasmid persistence

To get insight into the evolutionary trajectories by which plasmid pBP136Km and its coevolved MR-1 hosts adapted to each other, we focused on three clones (plasmid-host pair) each from lineages L1 and L4. We first determined whether mutations in the plasmid, the host, or both were responsible for the observed plasmid stabilization. This was done by constructing every possible combination of ancestral and evolved hosts and plasmids and comparing their plasmid persistence dynamics (Fig. 1). The plasmid population dynamics model previously described in our previous studies ^29,34,35^ was used to determine if the persistence dynamics of two bacteria-plasmid pairs where similar or not (see Materials and Methods and SI). When the increase in plasmid persistence is the result of plasmid evolution only, the evolved plasmid should exhibit the same persistence dynamics in the ancestral and evolved host. This was observed for clone L4-3 (Fig. 1, bottom right panel: pink and orange lines coincide). However, in this particular case the persistence of the ancestral plasmid was higher in the evolved host than in the ancestral host, suggesting also host evolution. Although the plasmid stabilization could be explained by the plasmid evolution only, the host apparently also underwent some changes that improve plasmid persistence. In contrast, when the increase of plasmid persistence is the result of host evolution only, the plasmid persistence dynamics should be the same for the evolved host with either the ancestral or evolved plasmid. This was the case for clone L4-1 (Fig. 1, bottom left panel: pink and blue lines are identical). Finally, when plasmid persistence increases as a result of the evolution of both plasmid and host, the co-evolved host-plasmid pair should exhibit significantly higher plasmid persistence than any other plasmid-host combination. This was observed for the four remaining clones. Thus within a single well-mixed liquid culture (L4) at least three clones with different evolutionary trajectories that improved the plasmid persistence still co-existed after 1000 generations. This is indicative of clonal interference.

### Several distinct molecular mechanisms can improve plasmid persistence

To identify the mutations that may explain the different plasmid persistence dynamics, the genomes of the six evolved plasmid-host pairs clones described in Fig. 1 were fully sequenced, and the sequence reads were mapped to their respective ancestral sequence (Table S1). The plasmid mutations are detailed below and can be summarized as follows: (i) small duplications in the replication initiation protein (*rep*) gene *trfA* (evolved plasmids pL1-1, pL1-2, pL1-3, pL4-2, pL4-3), (ii) acquisition of a transposon from the native plasmid(s) (pL4-3), and (iii) no genetic changes at all (pL4-1) (Fig. 2, Table S1).

**Figure 2.**
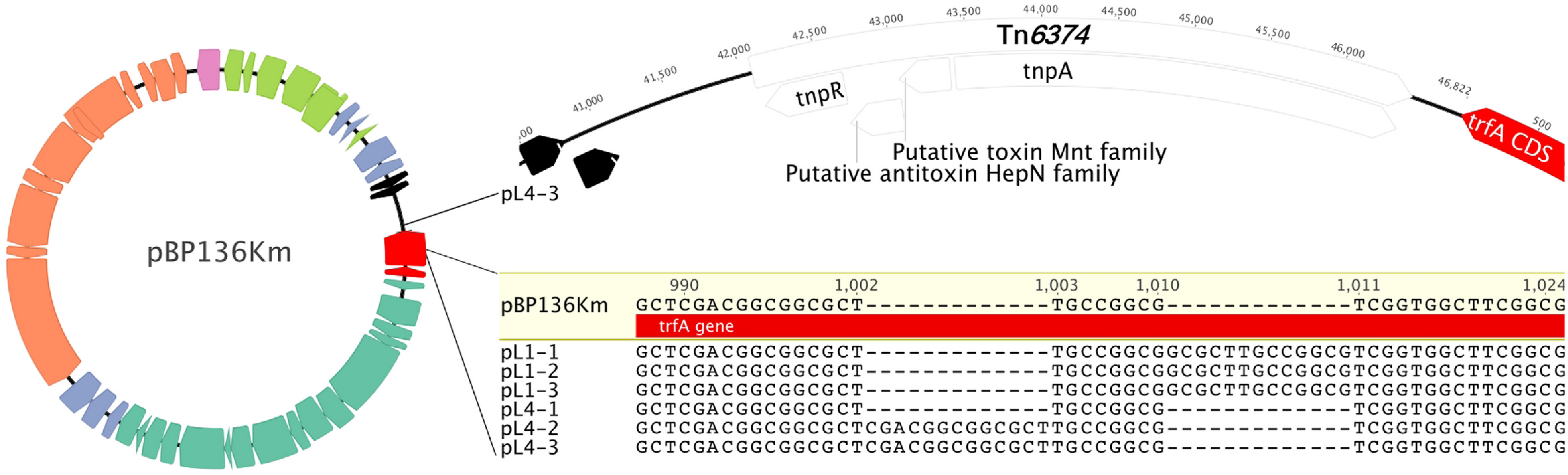
Map of plasmid pBP136Km, used in this study. Genes are highlighted by colors linked to their function: orange, conjugation; darker green, mating pair formation; red, replication; light blue, stable inheritance and partitioning; light green, regulation of plasmid functions; black, unknown function; pink, exogenous origin (inserted km resistance cassette). Top right shows the inserted transposon segment and its insertion location; bottom right shows the alignment of the *5’-trfA* region of the six plasmids to that of the ancestral plasmid.

First, all but one clone (L4-1) underwent one of two 13-bp duplication events in the 5’-end of the plasmid *trfA* gene (Fig. 2). These duplications were close to each other and they both introduced a frameshift leading to the same premature stop codon. As a result, TrfA1 was truncated but TrfA2 expression should be unaffected as the *trfA1* mutation was upstream of the start codon of TrfA2. In most IncP-1 plasmids, there are two versions of the Rep protein encoded by overlapping open reading frames: a long one (TrfA1) and a shorter one (TrfA2) with its one translational start codon downstream ^36^. In addition, plasmid pL4-3 had acquired a 4.4-kb segment in the *oriV-trfA* region. This fragment was identified as a transposon named Tn*6374*, which was present in the endogenous plasmids of MR-1 (Fig. 2). *In silico* analysis of the gene content of the transposon suggested it encodes a putative toxin-antitoxin (TA) system composed of two putative proteins sharing identities with the bifunctional TA system MNT/HEPN ^37,38^. Strikingly, the corresponding evolved clone L4-3 was the only one where plasmid mutations explained all of the improved plasmid persistence (Fig. 1, bottom right). Despite the significant effect of the *trfA* mutations on plasmid persistence in the other four clones (orange dashed lines in Fig. 1), they did not fully explain the great improvement in the remaining, suggesting additional mutation(s) in the host were required. Finally, no mutations were observed in the plasmid pL4-1 from clone L4-1, which is consistent with the finding that this host retained the ancestral plasmid as well as pL4-1.

To explain improved persistence of the plasmid in the clone L4-1 that didn’t show any plasmid mutations, we compared the chromosome sequence of this evolved host with the ancestor and found four SNPs (Fig. 3, Table S1). Two of these were detected in all the other clones from the same lineage, suggesting they were not responsible for the specific phenotype observed. However, the other two SNPs were unique to this clone; one was located in an intergenic space distant from any coding region, and the other one was in the *fur* gene, encoding the Fur protein, a well-known global transcriptional regulator protein ubiquitous in Gram-negative bacteria (Fig. 3) (Table S1) ^39,40^. The base substitution in the *fur* gene resulted in a change in amino acid sequence (A53 T) located in the DNA binding alpha-helix of the Fur protein ^41^. This change replaced a small non-polar amino acid with a polar amino acid, known to have major effects on alpha-helix structure. The influence of this amino acid substitution was predicted using the prediction tools PROVEAN (http://provean.jcvi.org/index.php) and SIFT (http://sift.bii.a-star.edu.sg/). Both models predicted this mutation would have a significant deleterious effect on the biological function of the protein. Thus in this evolved host the Fur protein function is most likely negatively affected, if not inactivated, suggesting that lowering or abolishing Fur function promotes the persistence of pBP136Km.

**Figure 3.**
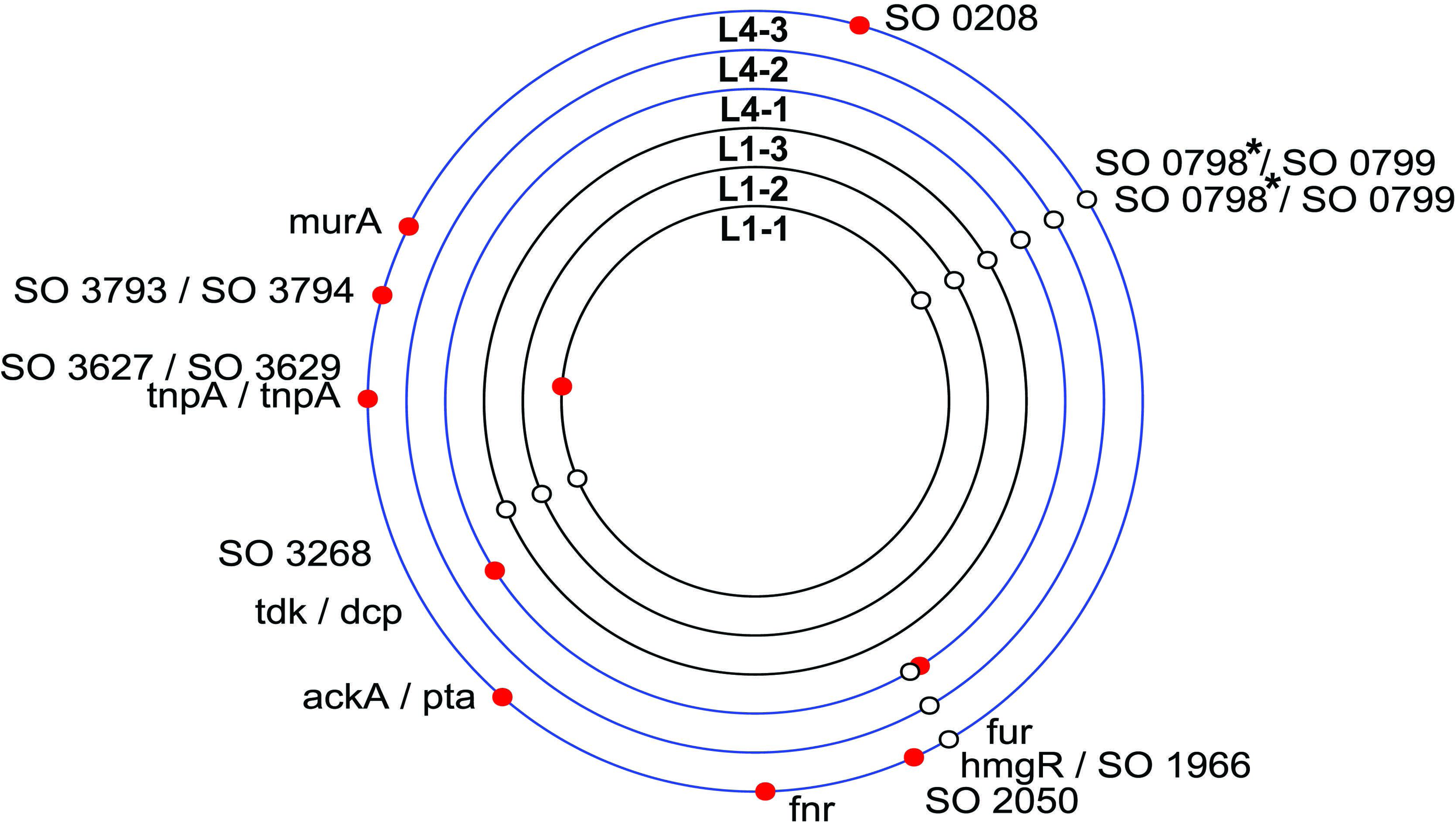
Identification of the chromosomal mutations in six evolved MR-1 strains. The mutations represented by circles are plotted on the circular chromosome of each isolate. A red circle denotes a mutation that is unique to the isolate. *: promoter under Fur regulation.

It is also noteworthy that in both lineages L1 and L4 two different SNPs arose upstream of the operon composed by SO_0798 and SO_0797, suggesting strong selection (Fig. 3). They encode, respectively, a TonB-dependent receptor and a periplasmic thioredoxin-family protein. It is striking however that transcription of these two CDS has been shown to be under negative control of the Fur protein^42^, plus the two SNPs occurred in the predicted “Fur box”, binding region of the Fur protein resulting in the repression genes expressions. These mutations could represent either adaptation to the environment or specific adaptation to the plasmid.

### The host’s Fur protein is involved in plasmid persistence

The effect of the Fur protein on plasmid persistence was evaluated by determining the persistence of ancestral pBP136Km in an existing MR-1 Δ*fur* mutant ^39^. In this mutant the segment within the *fur* gene that codes for the predicted DNA-binding domain and both metal-binding sites has been deleted. The ancestral plasmid was significantly more persistent in the Δ*fur* mutant than in wildtype MR-1, but not as persistent as in the evolved clone L4-1 (Fig. 4). This suggests that the Fur protein played at least a partial role in destabilizing a broad-host-range self-transmissible plasmid.

**Figure 4.**
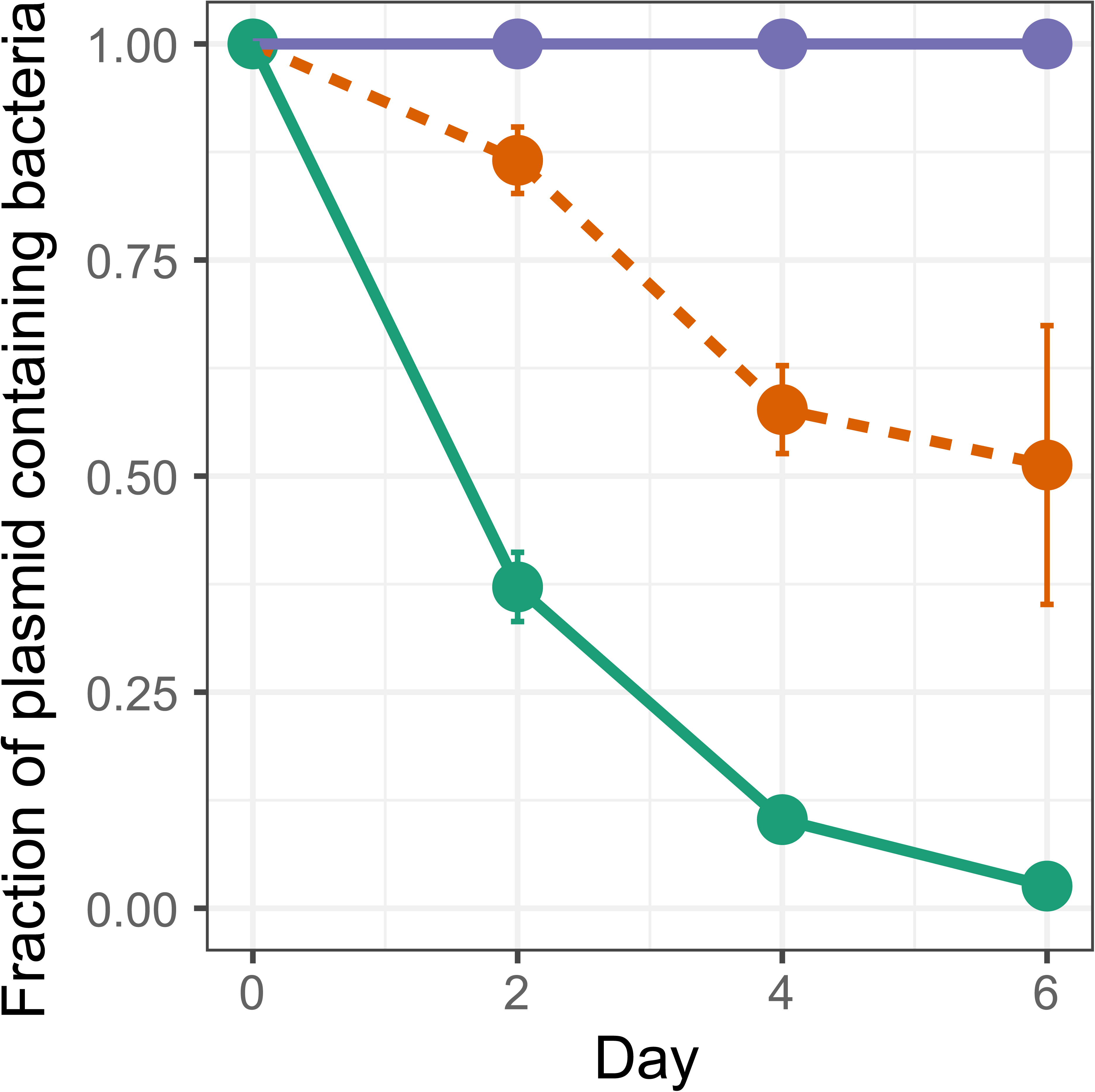
Persistence of the ancestral plasmid pBP136Km in the wildtype MR-1 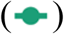 and in the *Afur* mutant 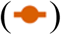 compared to the evolved clone L4-1 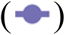. Each point is the mean of 3 individual replicates and the error bars show the standard deviation. The plasmid population dynamics model previously described^29^,^34^,^35^ was used to determine if the persistence dynamics of two bacteria-plasmid pairs where similar or not (see Materials and Methods and SI).

### Plasmid-host coevolution results in a shift in host range

Mutations in the 5’ region of *trfA1* have previously been shown to result in a shift in host range of the pBP136 mini-replicon ^26,43^. To determine whether the mutation observed here had a similar effect in the native plasmid pBP136Km, the persistence of the evolved plasmid pL1-1 was compared to that of the ancestral plasmid in the same hosts that were previously tested, *Pseudomonas putida* KT2440 ^33^ and *Sphingobium japonicum* UT26. The 13-bp duplication in the *trfA1* gene of this plasmid was identical to one previously described in the mini-replicon. As shown in Fig. 5, its presence significantly improved the persistence of the plasmid in both hosts, confirming that it can affect the plasmid’s long-term host range.

**Figure 5.**
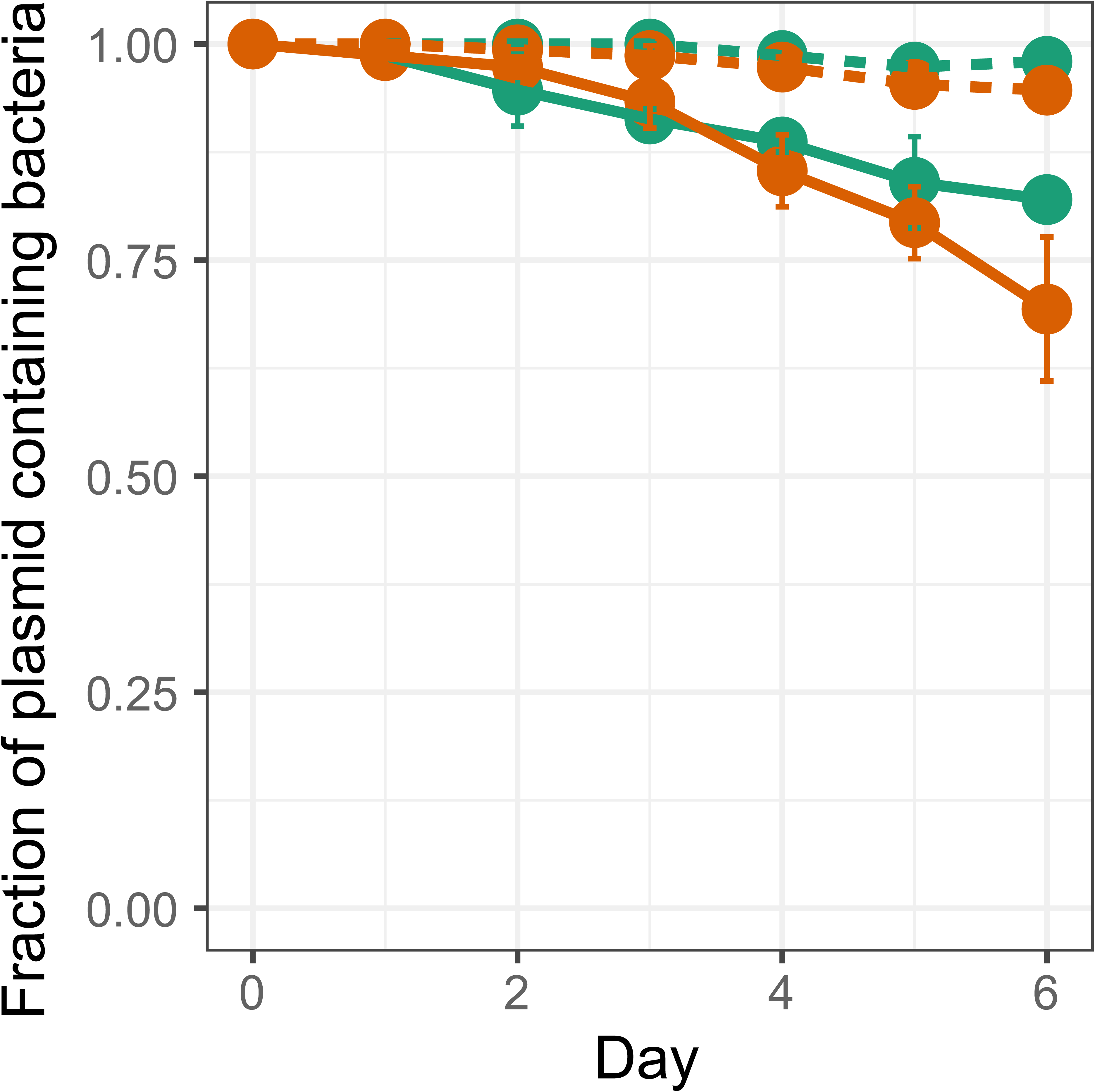
Persistence of ancestral (**—**) and evolved pL1_1 plasmid (––) in the naïve hosts *P. putida* 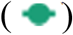 and *S. japonicum* 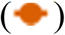. Data presented for *P. putida* have been previously published as supplementary information in Yano *et al,* 2016 ^33^. Each point is the mean of 3 individual replicates and the error bars show the standard deviation. The plasmid population dynamics model previously described^29,34,35^ was used to determine if the persistence dynamics of two bacteria-plasmid pairs where similar or not (see Materials and Methods and SI)

### The reported endogenous megaplasmid sequence of *Shewanella oneidensis* MR-1 in fact represents two independently replicating plasmids

Since in one of our clones the insertion of a transposon with a putative TA system in plasmid pBP136Km increased the plasmid’s persistence, we examined the transposon sequence and its flanking regions in the host genome. Originally, one endogenous plasmid of 161,613 base pairs had been described in strain MR-1 (accession number NC_004349.1) ^44,45^. Sequence analysis of that plasmid unraveled the presence of two identical copies of a transposon in the same orientation. This transposon contains a transposase closely related to the DDE transposase of the Tn*3* family (96% amino acid identity to the Tn*3* transposase of *Aeromonas salmonicida*, accession number WP042469084), and a site-specific serine recombinase (94% amino acid identity to the resolvase of the transposon in *Aeromonas salmonicida*, accession number WP017412788). As most Tn*3* family transposons display transposition immunity, resulting in the presence of only one copy per replicon ^46^, the presence of two copies in the same plasmid is very unlikely. In addition, the resolvase, encoded by the transposon, mediates recombination between directly repeated copies of a resolution site within the transposon region, splitting a cointegrate into two molecules, each carrying a single copy of the transposon ^47^. Thus, the arrangement of two transposon copies in the same orientation would be unusual and is expected to be unstable. Finally, agarose gel electrophoresis of entire plasmid extracts of our MR-1 strain during this study always showed the presence of two endogenous plasmids (Fig. S2, SI).

To verify that the previously reported mega plasmid is actually composed of two independent replicons, each one carrying a single copy of the transposon, the presence and the location of the transposon was characterized by PCR amplification across the putative transposon-plasmid junctions, followed by sequencing of the amplicons (Fig. S1 and Table S2, SI). The boundaries of the transposon were different than those of the NCBI sequence and suggested that the transposon is present as a single copy on each of the two circular replicons (Table S3, SI). For all these reasons we propose that the large plasmid sequence reported in the MR-1 Genbank sequence is in fact incorrect and should be split into two independent replicons of 142,357 bp and 19,256 bp, each containing a copy of Tn*3* family transposon Tn*6374* that transposed into our plasmid pBP136Km.

## DISCUSSION

Plasmid spread and persistence in bacterial communities is of great concern to human health because its contribution to the rapid spread of antibiotic resistance to human pathogens ^11,48^. To combat the spread of multidrug resistance it is therefore essential to understand the general mechanisms that allow a newly formed bacterial host-plasmid pair to (co)-evolve towards improved plasmid persistence in the absence of antibiotics ^49^. An experimental evolution approach has begun to unravel various adaptive mechanisms that drive plasmid-host co-adaption, but clear patterns have yet to emerge ^16,18,19,22,23,25–30^. This diversity of genetic solutions to improving plasmid persistence suggests that there may not be a general mechanism that could be targeted in future alternative antimicrobial therapies. However, in this study three adaptive trajectories shown to stabilize a conjugative drug resistance plasmid have been observed in previous studies, suggesting they are common: (i) mutations linked to DNA replication of plasmid and host which compensate the interference cost of the plasmid ^26,31,33^; (ii) the acquisition of a native transposon that carries a TA system ^29^; (iii) a compensatory mutation in a global regulatory pathway of the host ^16^. Based on these and previous findings we suggest common patterns of plasmid-host coevolution that can stabilize antibiotic resistance in bacteria, as discussed below.

In addition to its relevance to the spread of antibiotic resistance, this study contributes to discussions of host-symbiont evolution in a parasitism-mutualism continuum ^16^, and how compensatory evolution could drive evolution toward mutualism. We found that the original parasitic plasmid-host association in the absence of antibiotics evolved toward a more mutualistic interaction after the pair was evolved under conditions where both partners required each other to survive, i.e., in the presence of antibiotics.

In all clones wherein evolution of plasmid pBP136Km was observed, the mutations in the plasmid’s replication initiation protein (Rep) protein TrfA1 were the same as those previously demonstrated to improve the persistence of its non-transmissible mini-replicon in the same host. Such mutations, resulting in truncation of TrfA1 and expression of only one of the two intact Rep proteins, TrfA2, have been shown to lower the plasmid cost by reducing the protein affinity for the host helicase DnaB ^33^. In addition, as confirmed here, these *trfA* mutation resulted in a host range shift. Mutations in the Rep protein thus appear to represent a general evolutionary trajectory in *S. oneidensis* MR-1 that leads to higher persistence of a promiscuous IncP-1 plasmid and a shift in plasmid host range. It emphasizes the importance of plasmid-helicase interactions as a general adaptive strategy for plasmid-host adaptation, and parallels other recent studies that point towards the involvement of helicase mutations^31^ and Loftie-Eaton (personal communication).

Unlike our previous studies, here the *trfA* mutations alone were not sufficient to explain the high persistence of the plasmid after evolution. The main difference with the former studies is the presence of the conjugative machinery on the large version of the pBP136Km plasmid. Horizontal transfer can be costly to the host ^21^. This has been recently pointed out by an experimental evolution study that showed that deletion of the conjugative machinery was beneficial for plasmid maintenance in a population ^30^. Here, the conjugation functions were not affected, but increase of plasmid persistence required either host mutation(s) or acquisition of a putative TA system in addition to the *trfA* mutation.

The acquisition from a native plasmid of a transposon that encodes a putative TA and resolvase was entirely parallel to how the mini-pBP136Km stabilized in a different genus, Pseudomonas ^29^. This suggests that this genetic solution is a much more important mechanism of antibiotic resistance stabilization than hitherto expected. Here the acquired Tn*6374* contained a putative TA system of the family HEPNMNT and a resolvase (TnpR). The addiction mechanism of a TA system acquired by the newly introduced plasmid likely favors its vertical transmission by inhibiting the growth of cells that lost the plasmid. The transposon-encoded resolvase may further improve plasmid partitioning through resolving plasmid multimers before cell division^29,50–52^. Multidrug resistance plasmids that acquire a transposon-encoded TA system could thus persist longer in the absence of antibiotics and therefore more likely transfer to new bacteria. This evolutionary plasmid stabilization mechanism needs to be taken into account when trying to slow down the rapid spread of drug resistance in bacterial pathogens.

In one of the evolved clones, a SNP mutation in the global transcriptional regulator Fur of host MR-1 improved plasmid persistence. The persistence level of the ancestral plasmid in an MR-1 *fur* deletion mutant was also improved over that in the ancestral host, but not to the same extent. The SNP located in the DNA binding domain might have just changed the activity of the protein, or its ability to recognize a certain promoter, and not abolished it. It is striking that two other mutations found in the six evolved clones were in the predicted binding region of the Fur transcriptional regulator (Fur box) of genes known to be negatively regulated by Fur. Together these results suggest that Fur had a negative effect on plasmid persistence in this host, and mutation to decrease its regulatory effect were selected for under antibiotic selection. This hypothesis is supported by data collected by Lang *et al*. (2015) ^53^, who studied the differential gene expression in MR-1 following acquisition of the A/C plasmid pAR060302. They found that 446 genes in the MR-1 chromosome were differentially transcribed, among them the *fur* gene, which was upregulated (personal communication). The Fur protein is a pleiotropic regulator involved in the regulation of expression of wide diversity of genes in Gram-negative bacteria ^39,40^. Recently, mutations in other chromosomal genes have been shown to increase plasmid persistence by reducing the fitness cost of the plasmid, including a gene encoding a different global regulator ^16,28^. More specifically, Harrison *et al*. (2015) showed that a plasmid imposed a high cost to the bacterial host by upregulating a large amount of chromosomal genes. A mutation in a well-known bacterial two-component global regulatory system (*gacA/gacS*) significantly compensated for this cost by downregulating these genes back to normal levels ^16^. How exactly a novel plasmid influences the host’s transcriptional network is still an open question, but compensatory mutations in global regulatory proteins seem to represent another common evolutionary trajectory by which a bacterial host can counteract the interference cost imposed by horizontally acquired DNA.

To conclude, this study illustrates the diversity of evolutionary solutions that stabilize plasmid-encoded antibiotic resistance in the absence of any drugs, even within a single bacterial population. This underlines that plasmid-bacteria pairs are evolving on a rugged adaptive landscape with multiple evolutionary trajectories, and that clonal interference plays an important role in plasmid-host coevolutionary dynamics ^27^. Nevertheless, some common patterns of plasmid-host adaptation seem to emerge such as (i) compensatory mutations in global regulatory pathways, (ii) mutations reducing fitness cost imposed by plasmid DNA replication machinery, and (iii) the acquisition of a copy of native transposon that carries a TA system. Novel therapies to combat the spread of antibiotic resistance in pathogens should focus on bacterial targets that are involved in these common evolutionary mechanisms.

## METHODS

### Bacterial strains, plasmids and media

The plasmid host in this study was the Gamma-proteobacterium *Shewanella oneidensis* MR-1 ATCC 700550, isolated from river sediment (in this study named briefly MR-1). It was shown to poorly maintain plasmid pBP136 in the absence of selective pressure (see Fig. 1)^54^. For the host range experiment *Sphingobium japonicum* UT26SR*, Pseudomonas putida* KT2440, *Escherichia coli* MG165*5* and *E. coli* EC100 were used ^55^. All except the *S. japonicum* strain were grown in LB (Lysogeny Broth) culture medium, and *S. japonicum* UT26SR was grown in 1/3-LB (3.33 g Bacto tryptone, 1.67 g yeast extract and 5g NaCl per liter). All cultures were grown overnight, or for 2 days for *S. japonicum*, at 30°C, and where required the media were solidified with 1.5% (wt/vol) agar and supplemented with 50 µg.ml^−1^ kanamycin (km50) and/or 50 µg.ml^−1^ rifampicin (rif50) (LB-km and LB-rif). The MR-1 *fur* mutant strain (Fur protein deletion) was a gift from Dr. Zhou Jizhong ^39^. All experiments were performed under at least Biosafety Level 1 conditions in a Biosafety Level 2 laboratory.

Plasmid pBP136 is a cryptic IncP-1β plasmid isolated from *Bordetella pertussis* ^32^. It was found in a *Bordetella pertussis* strain isolated from an infant that died of whooping cough. Since this plasmid contained no selectable markers, plasmid pBP136Km was used here, a derivative of the wild-type plasmid containing a kanamycin resistance gene ^32^. Plasmids were transferred into new strains either by electroporation or by conjugation using a biparental mating method. Plasmids were isolated by a modified alkaline lysis protocol as described in Sambrook and Russell (2001)^56^ or by using the Qiagen^®^ Plasmid Midi Isolation Kit (Qiagen, Valencia, CA, USA).

### Evolution experiment

Plasmid pBP136Km was transformed by electroporation into an LB-adapted MR-1 strain ^26^. Five separate transformants were selected from a LB-km plate and transferred to five separate test tubes containing 5ml LB-km and incubated overnight at 30°C in a rotary shaker incubator (200 rpm); these represented the ancestors of five separately evolved populations called lineages L1-L5. Subsequently, each 24-h period (+/- 1 h), 4.9 µl of culture was transferred to new tubes containing 5mL LB-km, which were again incubated for 24 h, resulting in approximately 10 generations of growth per day. This was repeated daily for 100 days, or 1,000 generations. Cultures were archived in glycerol at −70 °C every 10 days. At generation 1,000 subsamples of each population were diluted and plated on LB-km agar and incubated overnight at 30°C. Several colonies were randomly selected from each lineage and archived in glycerol at −70°C. These clones were numbered and named after the lineages they were isolated from, thus clone L1-1 was isolate 1 from lineage 1. Plasmids isolated from these clones were named according to the isolate, thus pL1-1 is the plasmid found in clone L1-1.

### Plasmid persistence assays

Plasmid persistence was determined for 6 days as previously described ^57^. Briefly, triplicate cultures were grown overnight in LB-km and aliquots (4.9 µl) were transferred into 5 ml of fresh culture medium without antibiotic. The latter was repeated daily for 6 days. Each overnight culture was serially diluted and plated on LB agar plates. After overnight incubation at 30°C, 52 single colonies were randomly selected and replica plated on LB and LB-km agar. The ratio of the number of colonies grown on LB-km and LB was calculated and represents the fraction of plasmid-bearing cells. For *S. japonicum,* the protocol was the same except samples were transferred and plated every 2 days, and the medium used was 1/3 LB.

### Statistical analysis

Comparison of the dynamics of the plasmid persistence was carried out using a plasmid population dynamics model 34,35, as described by us previously ^29^. A more thorough description of the model and statistical methods is provided in the SI. Briefly, the Bayesian Information Criterion (BIC) was used to determine if the persistence dynamics of two bacteria-plasmid pairs where similar or not. Briefly, the model describes the dynamics of plasmid loss in a population using a set of underlying parameters: the frequency of plasmid loss, the fitness cost of plasmid carriage, and the conjugation frequency. By fitting the model to the data, we then determined if one or two sets of parameters more likely explained the two plasmid persistence dynamics that were compared with each other. When two sets were more likely to explain each of the dynamics separately, the strains being compared were considered to show a difference in plasmid persistence.

### DNA sequence analysis of ancestral and evolved strains

Total genomic DNA was extracted from 1mL of overnight culture of bacterial clones by using the GenElute^TM^ Bacterial Genomic DNA kit (Sigma-Aldrich, St. Louis, MO, USA). The quality and integrity of the gDNA was assessed on a 1% agarose gel and the concentrations were determined fluorometrically using Quant-iT™ PicoGreen^®^ dsDNA Reagent (ThermoFisher Scientific, Waltham, MA, USA) on a SpectraMax^®^ Paradigm^®^ Multi-Mode Microtiter Plate Reader (Molecular Devices, Sunnyvale, CA, USA). Full genome sequencing was performed by the IBEST Genomic Resources Core using a MiSeq sequencer (Illumina, San Diego, CA, USA) and 300 bp Paired-End Sample Preparation Kits (Illumina, San Diego, CA, USA). Prior to analysis the reads were preprocessed through a read-cleaning pipeline consisting of the following steps: 1) duplicate read pairs (possibly resulting from multi-cycle PCR reactions carried out as part of library preparation) were removed using a custom Python script; 2) sequences were then cleaned to remove sequencing adapters and low quality bases using the software package Seqyclean (https://github.com/ibest/seqyclean).

Identification of the mutations was performed with the open source computational pipeline breseq 0.25d (http://barricklab.org/breseq). Briefly, for each sample the cleaned reads were mapped to the reference sequences of each replicon: (1) pBP136Km, obtained by *in silico* introduction of the km cassette into the original pBP136 sequence (NC_008459) as previously described ^32^, (2) the *S. oneidensis* MR-1 chromosome (NC_004347.2) and (3) its native plasmid as assembled in GenBank (NC_004347.2), and the two putative native plasmids, here designated pSMR-1 and pLMR-1. All SNPs identified in the two ancestor clones that were sequenced were subtracted from the sequences of the evolved isolates. In addition, all suggested new junctions between disjoint regions of the reference sequence but where the scoring system of the pipeline classified them as ‘non assigned-junction evidence’ sequences were manually screened to verify their validity. A full description of the mutations that were identified is presented in Table S1, SI.

All possible mutations (point mutations, insertions or deletions) in pBP136Km were verified by designing primers in the flanking regions of the target, generating PCR amplicons using Phusion^®^ High-Fidelity DNA polymerase (New England Biolabs, Ipswich, MA, USA), and determining their sequence by Sanger sequencing (Elim Biopharm, Hayward, CA). The same protocol has been used to determine the location of the insertion of the transposon into pBP136Km, and to confirm the transposon-containing region of the native plasmids. All Sanger sequence manipulations (e.g., Sanger sequence analysis) were conducted in Geneious R7 (Biomatters, http://www.geneious.com).

## Data availability

All sequencing data pertaining to this project have been made available at the National Center for Biotechnology Information (SRA accession number SRP105365). Other raw data are available upon request.

## ACKNOWLEDGEMENTS

This work was supported by the National Institute of Allergy and Infectious Diseases grant R01 AI084918 of the National Institutes of Health (NIH). The genome sequencing was carried out by the IBEST Genomics Research Core, and made possible thanks to NIH National Institute of General Medical Sciences under Award Number P30 GM103324. Thibault Stalder was funded for this work by the National Science Foundation BEACON Center for the Study of Evolution in Action under cooperative agreement no. DBI-0939454. Chris Renfrow obtained a student undergraduate research fellowship through the National Science Foundation UBM grant, award number DUE 1029485 (UI-WSU Program in Undergraduate Mathematics and Biology). We thank Dr. Zhou Jizhong who kindly provided the *Shewanella oneidensis* MR-1 *fur* mutant, and Dr. Jose Ponciano for his support on the statistical analysis using the plasmid population dynamics model.

## CONTRIBUTIONS

EMT conceived the project; TS and EMT wrote the manuscript, LMR helped in the writing of the manuscript; TS, LMR, CR, ZM performed experiments; TS and LMR analyzed the data, LMR and HY resolved the 2 endogenous plasmids.

